# Integrative Network Analysis Reveals Organizational Principles of the Endocannabinoid System

**DOI:** 10.1101/2025.10.28.685143

**Authors:** Aanya Shridhar, Sugyan Mani Dixit, Anthony Torres, Reggie Gaudino

## Abstract

**Background:** The endocannabinoid system (ECS) is a complex signaling network that regulates diverse physiological processes, including pain, mood, metabolism, and immune response, through coordinated interactions among receptors, enzymes, and lipid-derived ligands. Despite extensive research on individual ECS components, the systems-level organization and network resilience of the ECS remain underexplored. Here, we present a systems-level analysis of the ECS that integrates protein–protein and protein–chemical interactions into a unified network framework.

**Methods:** We constructed integrated ECS networks that combine protein–protein and protein–chemical interactions, utilizing data from multiple public databases. Network analyses were performed in Python using NetworkX to assess molecular connectivity and interaction topology. We utilized centrality measures to identify major hubs, employed community detection algorithms to examine the clustering of nodes, and performed targeted perturbations by sequentially removing the top-ranked nodes based on degree and betweenness centrality to assess network robustness.

**Results:** Centrality analyses identified the primary cannabinoid receptors, cannabinoid receptor 1 (CNR1) and cannabinoid receptor 2 (CNR2), as major hubs with extensive connectivity to endogenous and exogenous ligands. Non-canonical receptors, including transient receptor potential vanilloid 1 (TRPV1) and G-protein coupled receptor 55 (GPR55), also emerged as highly ranked nodes across multiple centrality measures, underscoring their integrative roles within the ECS signaling pathway. Community detection revealed biologically meaningful modules centered around receptor and metabolic clusters, with CNR1, CNR2, anandamide (AEA), 2-arachidonoylglycerol (2-AG), and major phytocannabinoids maintaining key network connectivity. Perturbation analyses demonstrated that removal of top hubs, particularly CNR1, caused pronounced losses in edge connectivity and disrupted signaling pathways among cannabinoids. However, the redistribution of influence toward CNR2 and GPR55 under multi-node removal conditions revealed compensatory plasticity and resilience within the ECS network.

**Conclusion:** This systems-level study highlights the hierarchical and robust architecture of the ECS. The identification of hub nodes, functional communities, and compensatory mechanisms provides insight into how the ECS maintains signaling integrity in the face of perturbation. These findings establish a network-based framework for studying cannabinoid biology and may inform future therapeutic strategies targeting the ECS and its interacting molecular pathways.

## Background

The endocannabinoid system (ECS) is a multifaceted regulatory network that maintains physiological homeostasis across diverse organ systems [1–3]. It is composed of endogenous ligands known as endocannabinoids, their cognate receptors, and the enzymes that mediate their synthesis and degradation [1,4]. Endocannabinoids are cannabinoids produced within the human body, in contrast to phytocannabinoids, which are derived from a number of plants, of which the best known is *Cannabis sativa* [5–7], and synthetic cannabinoids, which are laboratory-engineered compounds designed to mimic or modulate the effects of natural cannabinoids for research and therapeutic applications [1,4]. Through the coordinated activity of these molecular components, the ECS influences a broad spectrum of biological processes, including pain perception, inflammation, memory, and mood regulation [8–10].

Two primary receptors define much of the functional landscape of the ECS. Cannabinoid receptor 1 (CB1, also called CNR1) is predominantly expressed in the brain and central nervous system, where it regulates mood, appetite, pain perception, and cognitive processes [1,8–10]. Cannabinoid receptor 2 (CB2, also called CNR2) is primarily located in immune cells and peripheral tissues, where it regulates inflammatory and immune responses [1,8–10]. The ECS also interfaces with intracellular kinase signaling pathways that control cell survival, synaptic plasticity, and metabolic regulation. This extensive interconnectedness underscores how the ECS integrates with broader cellular signaling networks [1–4,8–12]. When the processes governing endocannabinoid synthesis and degradation become dysregulated, the resulting imbalance in “endocannabinoid tone” has been associated with a range of pathological conditions, including chronic pain, neurodegenerative disorders, metabolic syndromes, and cancer [11–15].

The growing acknowledgment of the ECS’s role in human physiology has made it a key target for both basic and clinical research, particularly in understanding how endogenous and exogenous compounds can modulate it’s signaling [16–19]. The active compounds delta-9-tetrahydrocannabinol (Δ9-THC) and cannabidiol (CBD), derived from medical cannabis, interact with and modulate the activity of cannabinoid receptors, specifically CNR1 and CNR2, influencing endocannabinoid signaling and therapeutic outcomes [20–22]. Specifically, CBD-rich formulations with minimal THC content are increasingly studied and prescribed for conditions such as epilepsy, anxiety, and inflammation [20–22]. Despite this rising interest, the ECS remains insufficiently characterized as a networked system, particularly in terms of how its molecular components interact and respond to changes [1]. A systems-level understanding of these interactions is critical for identifying novel therapeutic targets and developing strategies that more precisely modulate ECS function [23].

In this study, we employed a systems biology framework to characterize the ECS as an integrated molecular network. We constructed interaction networks encompassing protein–protein and protein–chemical relationships within the system. Through network analysis, we identified key nodes using multiple centrality measures and performed clustering analyses to delineate groups of molecules with shared functional characteristics. To further examine network dynamics, we introduced perturbations to assess robustness and to explore how the ECS responds to simulated stress. Collectively, this framework provides a systems-level representation of the ECS, offering insight into its organization, stability, and potential applications in therapeutic development.

## Methods

### Data Collection and Network Construction

To construct a comprehensive representation of the ECS, we compiled proteins, chemicals, protein–protein interactions (PPIs), and protein–chemical interactions (PCIs) from multiple databases (Figure 1). In the resulting networks, nodes represent ECS proteins or chemicals, and edges reflect experimentally supported interactions. All entities were annotated with structural and functional metadata to facilitate reproducibility and public access.

**Figure 1:**
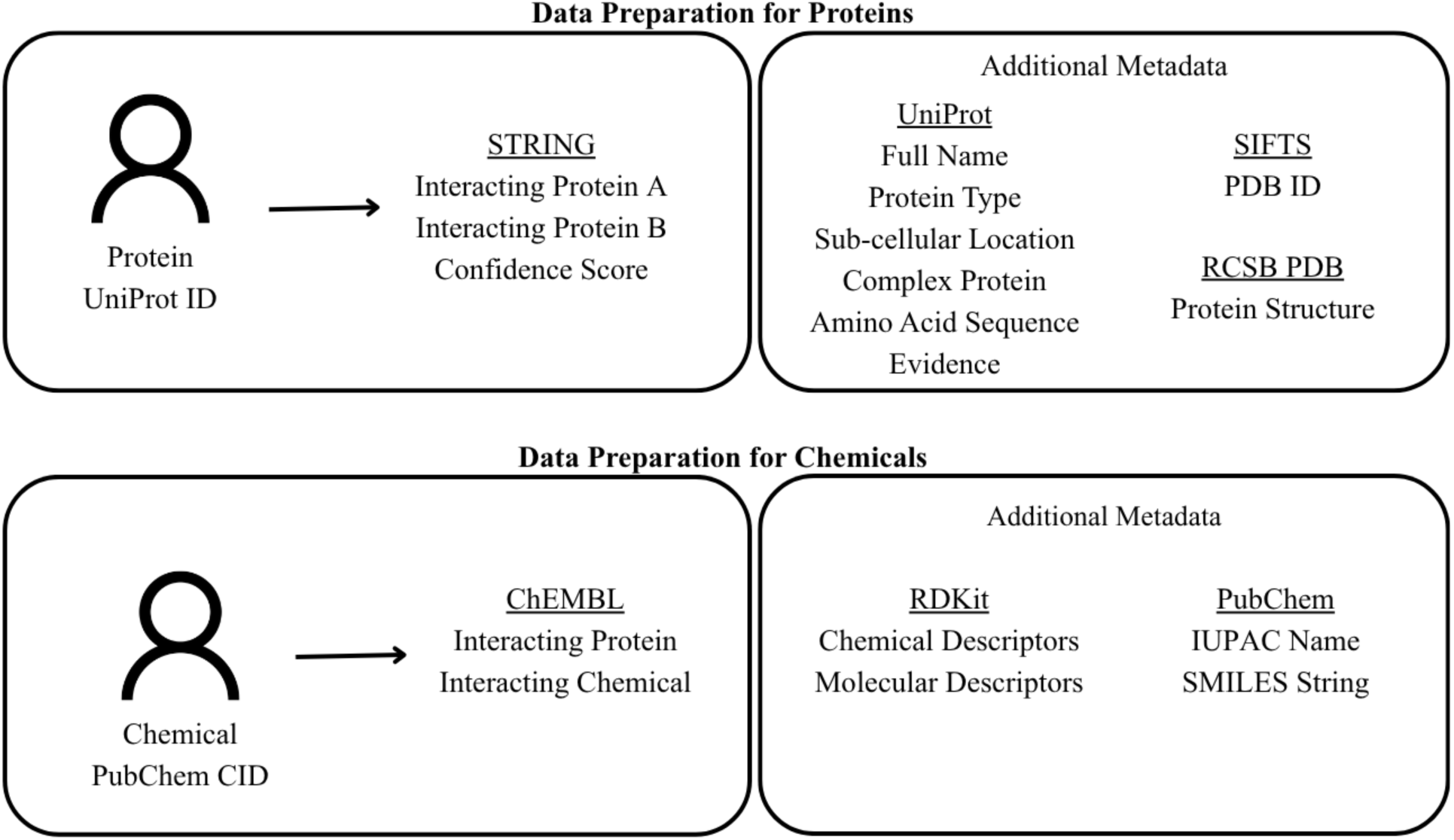
Data preparation pipeline for proteins and chemicals. Schematic depiction of the workflow used to compile protein–level and chemical–level data for network construction. Protein nodes (UniProt IDs) were annotated with interaction data from STRING, including interacting pairs and confidence scores. Additional metadata were integrated from UniProt, SIFTS, and RSCB PDB. Chemical nodes (PubChem CIDs) were annotated with interaction data from ChEMBL. Additional metadata were pulled from PubChem and calculated using RDKit.

#### Proteins

A curated set of forty ECS-associated proteins, including receptors CNR1 and CNR2, enzymes fatty acid amide hydrolase (FAAH), monoacylglycerol lipase (MGLL), and N-acyl-phosphatidylethanolamine phospholipase D (NAPEPLD), and signaling partners, was compiled from prior literature [23] and expanded using UniProt and STRING database queries [24,25]. Core attributes, including amino acid sequence, sub-cellular localization, and complex membership, were retrieved from UniProt [24]. Structural mappings to the Protein Data Bank were obtained via SIFTS, and protein structures were extracted from RCSB PDB [26,27]. Protein–protein interactions were collected from STRING using a confidence threshold of 0.7 to reduce false positives [25]. STRING confidence scores integrate multiple evidence types to estimate the likelihood of a true interaction, with values approaching one indicating higher confidence. All protein entries and protein-protein interactions were derived from human (*Homo sapiens*) data.

#### Chemicals

Forty ECS-relevant chemicals, including endocannabinoids anandamide (AEA) and 2-arachidonoylglycerol (2-AG), phytocannabinoids Δ9-THC and CBD, and selected eicosanoids, were curated from prior literature [23] and expanded using PubChem and ChEMBL database queries [28,29]. Chemical metadata, including IUPAC names and SMILES strings, were retrieved from the PubChem database [28]. Molecular descriptors, including molecular weight, partial charges, and topological fingerprints, were calculated using RDKit to serve as metadata [30]. Protein–chemical interactions were obtained from ChEMBL and filtered to retain only interactions involving the predefined ECS proteins, ensuring network specificity [29].

### Network Construction and Analysis

Three networks were constructed using NetworkX: a protein–protein only network, a protein–chemical only network, and a combined network containing all interactions [31]. Network topology was quantified using centrality metrics, including betweenness, closeness, degree, and eigenvector. These measures identify nodes that are highly connected, influential, or serve as critical bridges within the ECS, providing insight into the system’s organizational hierarchy [32].

### Network Clustering

Functional modules within the ECS network were identified using the Louvain algorithm, which maximizes modularity by clustering nodes with preferential interactions [33]. This approach highlights groups of proteins and chemicals that are likely to share biological functions or participate in coordinated processes.

### Network Perturbations

Network robustness was evaluated by targeting the removal of specific nodes. Specifically, the nodes with the highest betweenness and degree centrality, namely the top one and top three nodes from each metric, were systematically removed to simulate the disruption of crucial, highly connected components. After each removal, the network was reassessed to determine the number of remaining interactions and connections and to evaluate changes in its overall structure. Centrality metrics were recalculated to identify shifts in the relative importance of the remaining nodes and to understand how the network reorganized in response to the loss of these critical elements [23,34,35].

### Code and Data Availability

All data and code used in this study are available in a GitHub repository at https://github.com/aanyashridhar/ECS-Network.

## Results

The combined ECS network, which integrates both protein–protein and protein–chemical interactions, comprised 80 nodes and 218 edges and served as the primary framework for subsequent analyses (Figure 2a, Tables S1-S4). In this network, proteins occupy the central core, while chemicals are positioned toward the periphery. This structural organization reflects the biological architecture of the ECS, where proteins act as molecular hubs mediating signaling and metabolic processes, and chemicals serve as ligands or substrates that connect through these central protein nodes. The resulting topology suggests that the ECS is organized around a dense protein interaction core that enables diverse chemical inputs to converge on shared molecular pathways.

**Figure 2:**
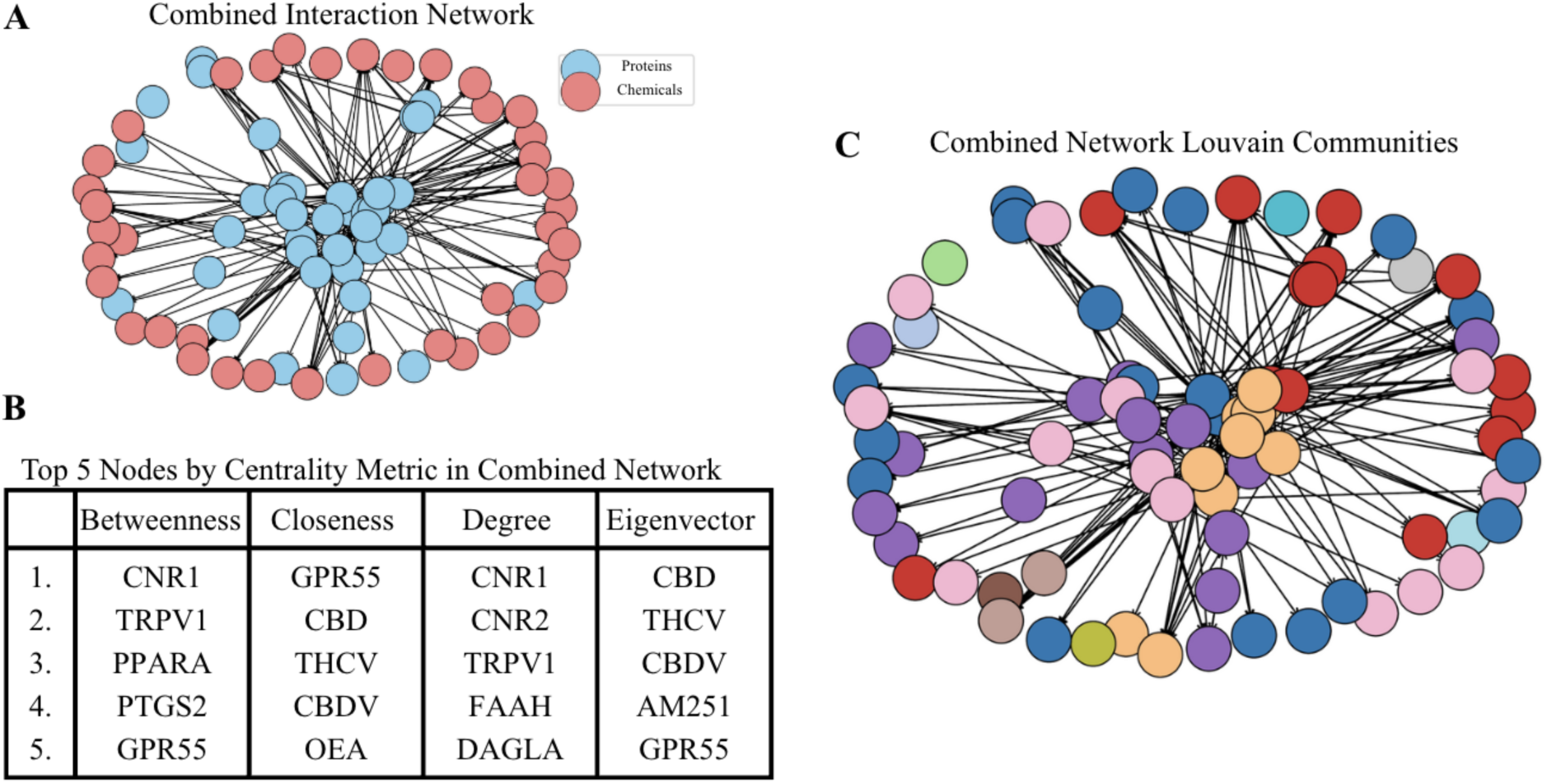
Combined interaction network and node characterization. **A)** Visualization of the integrated network showing proteins (blue) and chemicals (red) connected through documented interactions. The network has 80 nodes and 218 edges. **B)** Top five nodes ranked by centrality measures in the combined network. Results highlight distinct sets of influential nodes across betweenness, closeness, degree, and eigenvector centrality metrics, underscoring multiple dimensions of node importance. Full rankings and centrality metric calculations can be found at https://github.com/aanyashridhar/ECS-Network. **C)** Louvain community detection applied to the combined network, partitioning the nodes (denoted by distinct colors) into modular clusters based on interaction density, which highlights potential functional or mechanistic groupings within the ECS network.

For comparison, the protein–protein and protein–chemical subnetworks, analyzed separately, contained 40 nodes and 92 edges, and 80 nodes and 126 edges, respectively. These subnetworks are shown in Figures S1–S4 and Tables S1–S4, and provide additional insight into the distinct interaction patterns that contribute to the overall connectivity of the ECS.

### Network centrality analysis reveals the hierarchical organization of the ECS and provides insight into connectivity and influence patterns

To understand the structural organization of the ECS network, we quantified key centrality measures that reflect node influence and connectivity (Figure 2b, Tables S11-S12). The analysis revealed distinct patterns in how proteins and chemicals occupy topological positions within the combined network.

The five nodes with the highest betweenness centrality [32] in the combined network were CNR1, transient potential vanilloid receptor 1 (TRPV1), peroxisome proliferator-activiated receptor alpha (PPARA), cyclooxygenase-2 (PTGS2), and G-protein coupled receptor 55 (GPR55), in order of rank (Figure 2b). High betweenness centrality in these proteins indicates that they act as major conduits through which information and signaling flow across the ECS network, underscoring their integrative roles across chemical and protein subnetworks. As one of the two primary cannabinoid receptors, CNR1 connects to numerous cannabinoids and signaling components, serving as a central bridge in the ECS [16,36]. The second-highest ranked node, TRPV1, though not a canonical cannabinoid receptor, is a ligand-gated ion channel that mediates pain perception, inflammation, and thermoregulation [37]. It is strongly activated by cannabinoids, positioning it as a key integrative node within the ECS. The third-ranked PPARA is a nuclear receptor linking cannabinoid signaling to metabolic regulation [38]. The fourth-ranked enzyme, PTGS2, converts endocannabinoids into prostaglandins, thereby integrating the ECS with inflammatory and pain pathways [39]. Finally, GPR55, ranked fifth, is a G-protein coupled receptor (GPCR) that functions as a major non-cannabinoid receptor, further expanding the ECS signaling landscape [40].

The five nodes with the highest closeness centrality [32] in the combined network were GPR55, CBD, tetrahydrocannabivarin (THCV), cannabidivarin (CBDV), and oleoylethanolamide (OEA), in descending order (Figure 2b). High closeness centrality in these compounds suggests that they can efficiently reach many nodes in the network through relatively few interactions, consistent with their broad influence on ECS signaling dynamics. GPR55 interacts with multiple cannabinoids and receptors, resulting in short path lengths across the network [40]. CBD, THCV, and CBDV follow closely, consistent with their established promiscuity across diverse ECS targets, including CNR1, CNR2, TRPV channels, and GPCRs [20]. The fifth-ranked OEA, an endocannabinoid-like lipid, shares biosynthetic and signaling pathways with canonical endocannabinoids but does not bind directly to the cannabinoid receptors CNR1 or CNR2 [41].

Nodes ranked highest by degree centrality [32] in the combined network were CNR1, CNR2, TRPV1, FAAH, and diacylglycerol lipase alpha (DAGLA), in order (Figure 2b). Nodes with high degree centrality function as major hubs within the network, reflecting their broad engagement across ECS components and highlighting their central role in maintaining network connectivity. CNR1 and CNR2, the two canonical cannabinoid receptors, exhibit extensive connectivity across both protein and chemical domains [16,36]. TRPV1, again, emerges as a highly connected integrator of cannabinoid and non-cannabinoid signaling [37]. The fourth-ranked FAAH, a degradation enzyme [42], and the fifth-ranked DAGLA, a biosynthetic enzyme [43], emphasize the essential role of metabolic turnover in regulating ECS tone and signaling plasticity.

Finally, nodes ranked highest by eigenvector centrality [32] in the combined network were CBD, THCV, CBDV, AM251, and GPR55, in descending order (Figure 2b). High eigenvector centrality indicates that these nodes are not only well-connected themselves but also closely linked to other influential nodes, emphasizing their role in shaping highly interactive subnetworks within the ECS. The prominence of CBD, THCV, and CBDV highlights the influence of phytocannabinoids within the network, as they engage several central receptors and enzymes [20]. AM251, a synthetic CB1 antagonist, ranks fourth due to its strong association with the CNR1 receptor, reflecting its frequent use in receptor-targeting studies [44]. GPR55 reappears among the top five, reinforcing its status as a non-canonical yet highly connected receptor within the ECS [40].

### Network clustering analysis uncovers the modular architecture of the ECS and identifies biologically meaningful groupings of receptors, enzymes, and ligands

Clustering analysis identified thirteen distinct communities within the ECS network (Figure 2c, Tables S5-S6). The largest community consisted of six proteins and thirteen chemicals, while the smallest communities included only a single node (Tables S5-S6). Although these isolated nodes were formally classified as communities by the algorithm, they do not represent meaningful biological groupings. The overall modularity of 42.77% (Table S6) falls within the expected range for biological networks (typically 30–70%) [45], indicating a balance between connectivity and modular organization.

Among the identified clusters, two communities were particularly informative. The first and largest, designated community 0 (Table S5), captured the core signaling architecture of the ECS. This cluster included the two primary cannabinoid receptors (CNR1 [16,36] and CNR2 [36]), PLC enzymes (phospholipase C epsilon 1 (PLCE1) [46] and phospholipase C beta 1 (PLCB1) [46]), signaling partners (inhibitory Guanine nucleotide-binding protein G alpha subunits 1–3 (GNAI 1–3) [47]), numerous synthetic cannabinoids (WIN55,212-2 [48], WIN55,212-3 [48], JWH-018 [49], JWH-073 [49], JWH-133 [49], CP55,940 [50], SR141716A [51], SR144528 [51], and AM251 [44]), and endocannabinoid AEA [52,53]. The GNAIs represent canonical intracellular signaling mediators in the ECS, while synthetic cannabinoids and AEA illustrate the diversity of ligand–receptor interactions within the system. This cluster can be viewed as the cannabinoid receptor group, showing the cannabinoid receptors and their immediate effectors clustering strongly together. This organization reinforces the biological validity of the network and demonstrates that its modular structure captures known ECS signaling relationships.

The second notable cluster, community 2 (Table S5), which is the fifth largest community, centered on enzymes and molecules involved in the metabolism of endocannabinoids. This community contained degradation enzymes (FAAH [42], MGLL [54], abhydrolase domain-containing protein 6 (ABHD6) [55,56], and abhydrolase domain-containing protein 12 (ABHD12) [55,56]), synthesis enzymes (NAPEPLD [57], DAGLA [43], and diacylglycerol lipase beta (DAGLB) [43]), non-canonical receptor GPR55 [40,58], transporter (fatty acid-binding protein 7 (FABP7) [59]), and endocannabinoid 2-AG [53]. Together, these nodes represent the metabolic group of the ECS, highlighting the tightly coupled nature of endocannabinoid synthesis, degradation, and transport. The inclusion of GPR55 within this community further underscores the functional integration of metabolic and signaling processes that maintain ECS homeostasis.

### Network perturbation analysis illustrates how changes to key nodes reshape ECS connectivity, revealing dependencies within its signaling network

A total of four perturbations were performed for each network, including the removal of the top one node by betweenness centrality, the top three nodes by betweenness centrality, the top one node by degree centrality, and the top three nodes by degree centrality. The protein–chemical only network was limited to two perturbations because betweenness centrality relies on shortest path calculations between nodes, which are not well-defined in networks where chemicals only connect to proteins, allowing only degree-based removals. Across all networks, this approach produced a total of ten perturbations (Table 1). After each perturbation, the change in the number of edges before and after node removal was calculated as a high-level indicator of network impact. Perturbation by betweenness centrality in the protein–protein only network produced the most pronounced effect, with a 40.22% reduction in edge count. In general, removing the top one node, regardless of metric, decreased edge count by approximately 14–16%, whereas removing the top three nodes caused a larger reduction of roughly 34–40%. Notably, CNR1 was removed at least once in all networks, regardless of whether it was identified as a top-ranked node based on betweenness or degree centrality, underscoring its structural importance within the ECS.

**Table 1:** Network perturbation analysis across protein–protein, protein–chemical, and combined ECS networks. Summary of robustness simulations following targeted removal of highly central nodes (top one or top three rankings in betweenness or degree centrality). For each perturbation, the table shows the number of original nodes and edges, number of edges after perturbation, and percent change in edge count.

| Network | Perturbation Metric | Removal Type | Removed Node(s) | Original # Nodes | Original # Edges | Perturbed # Edges | % Change in Edges |
| --- | --- | --- | --- | --- | --- | --- | --- |
| Protein–Protein | Betweenness Centrality | Top 1 | CNR1 | 40 | 92 | 77 | 16.30 |
| Protein–Protein | Betweenness Centrality | Top 3 | CNR1, PPARA, PTGS2 | 40 | 92 | 55 | 40.22 |
| Protein–Protein | Degree Centrality | Top 1 | FAAH | 40 | 92 | 79 | 14.30 |
| Protein–Protein | Degree Centrality | Top 3 | FAAH, CNR1, GPR55 | 40 | 92 | 65 | 29.35 |
| Protein–Chemical | Degree Centrality | Top 1 | CNR1 | 80 | 126 | 105 | 16.67 |
| Protein–Chemical | Degree Centrality | Top 3 | CNR1, CNR2, TRPV1 | 80 | 126 | 82 | 34.92 |
| Combined | Betweenness Centrality | Top 1 | CNR1 | 80 | 218 | 184 | 15.60 |
| Combined | Betweenness Centrality | Top 3 | CNR1, TRPV1, PPARA | 80 | 218 | 141 | 35.32 |
| Combined | Degree Centrality | Top 1 | CNR1 | 80 | 218 | 184 | 15.60 |
| Combined | Degree Centrality | Top 3 | CNR1, CNR2, TRPV1 | 80 | 218 | 153 | 29.82 |

To evaluate how highly connected and influential nodes contribute to the stability of the ECS network, we performed targeted perturbations informed by network topology. Nodes were selected for removal based on their rankings in betweenness and degree centrality, which capture information flow and connectivity, respectively. For each perturbation, key centrality measures, including betweenness, closeness, degree, and eigenvector centrality, were recalculated to assess changes in network organization and influence distribution. The percent change in each metric was standardized relative to the distribution of changes across all nodes. Perturbations involving the removal of the top one or top three nodes by betweenness centrality (Figure 3) and degree centrality (Figure 4) were used to examine how hub disruption alters ECS structure and signaling balance.

**Figure 3:**
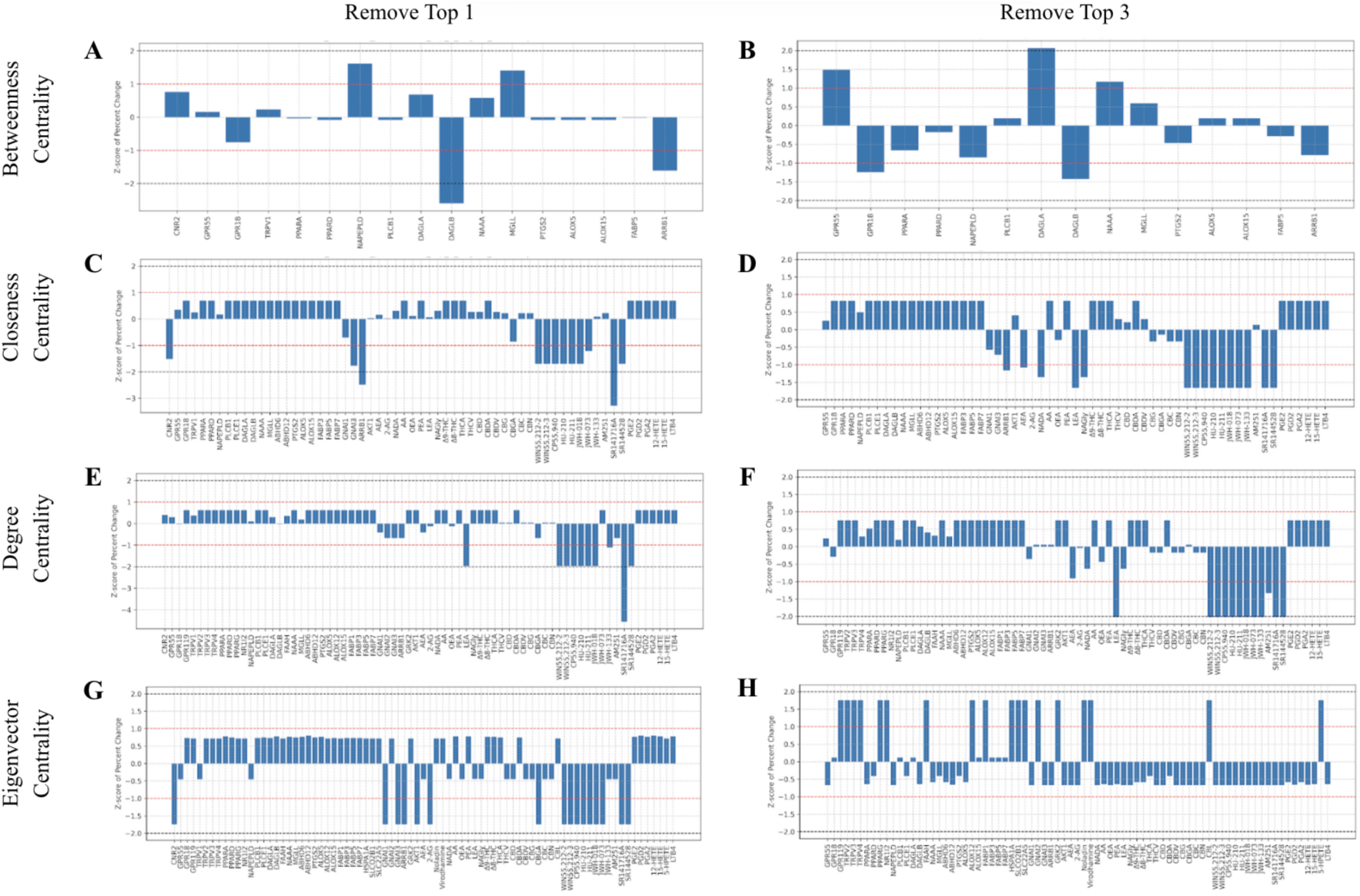
Network perturbation effects on node centrality following removal of high-betweenness nodes. Bar plots depict standardized changes (z-scores) in centrality metrics of the nodes after perturbation. The red and black horizontal dotted lines denote absolute z-scores of 1 and 2, respectively. **A-B)** Betweenness centrality recalculated after removal of the top one **(A)** and top three **(B)** nodes by betweenness. **C-D)** Closeness centrality recalculated after removal of the top one **(C)** and top three **(D)** nodes by betweenness. **E-F)** Degree centrality recalculated after removal of the top one **(E)** and top three **(F)** nodes by betweenness. **G-H)** Eigenvector centrality recalculated after removal of the top one **(G)** and top three **(H)** nodes by betweenness.

**Figure 4:**
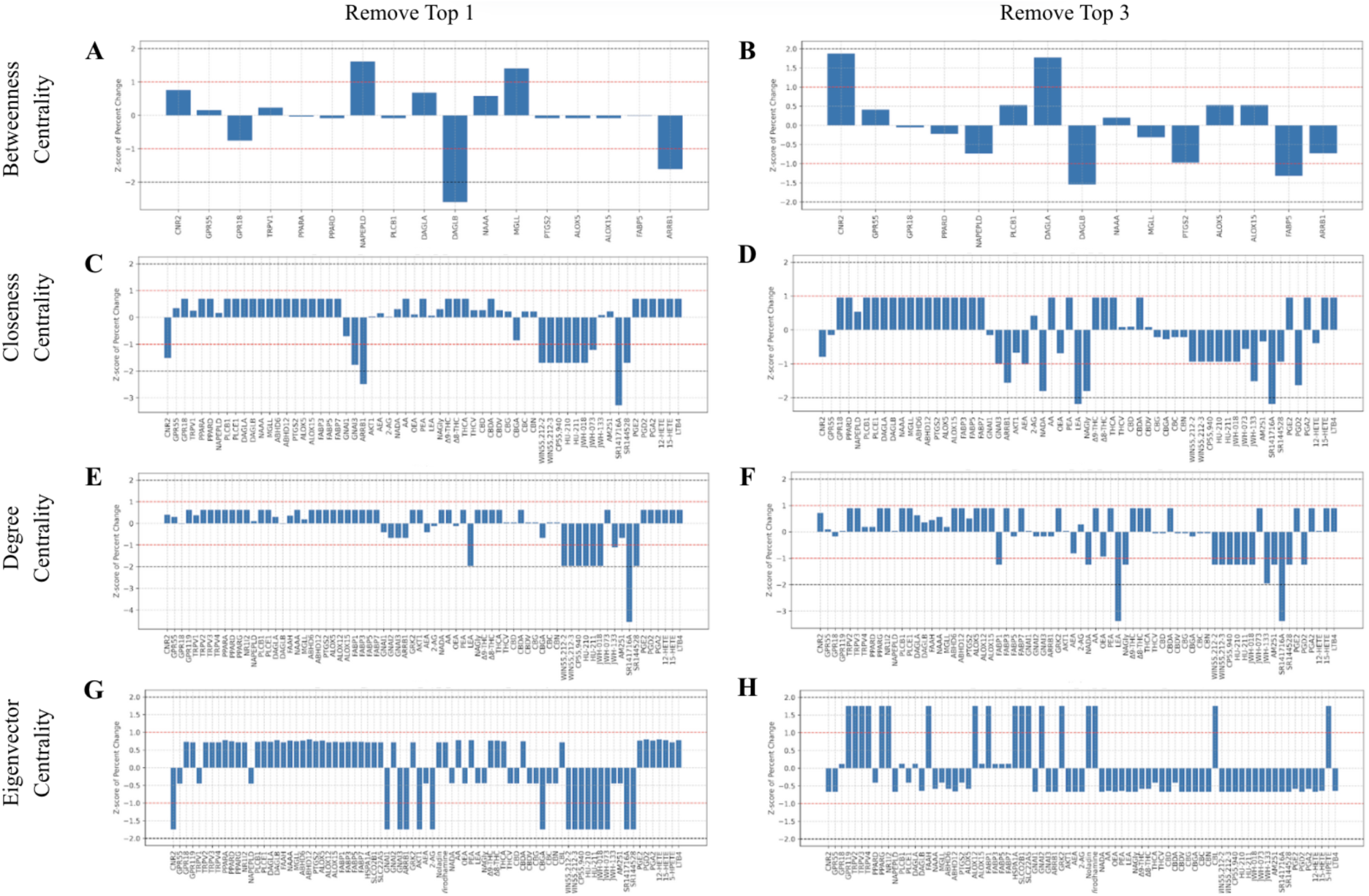
Network perturbation effects on node centrality following removal of high-degree nodes. Bar plots depict standardized changes (z-scores) in centrality metrics of the nodes after perturbation. The red and black horizontal dotted lines denote absolute z-scores of 1 and 2, respectively. **A-B)** Betweenness centrality recalculated after removal of the top one **(A)** and top three **(B)** nodes by degree. **C-D)** Closeness centrality recalculated after removal of the top one **(C)** and top three **(D)** nodes by degree. **E-F)** Degree centrality recalculated after removal of the top one **(E)** and top three **(F)** nodes by degree. **G-H)** Eigenvector centrality recalculated after removal of the top one **(G)** and top three **(H)** nodes by degree.

Perturbation analysis revealed that the removal of highly connected or central nodes caused pronounced shifts in network organization, highlighting which nodes and interactions are most critical for ECS function. In particular, in the absence of CNR1 [16,36], receptors CNR2 [36] and GPR55 [40] increased in centrality, reflecting their compensatory relevance within the network (Figures 3a, 3b, 4a, and 4b). CNR2 and GPR55 are important receptors that can partially maintain connectivity and signaling when CNR1 is absent, indicating redundancy and resilience in the ECS. Conversely, degradation enzymes such as DAGLB [43] showed decreases in betweenness centrality by over two standard deviations, suggesting that key signaling or metabolic nodes lose influence when primary receptors are disrupted (Figures 3a and 4a). Similarly, the endocannabinoid-like lipid linoleoyl ethanolamide (LEA) [41] exhibited a drop in closeness centrality by nearly two standard deviations following the removal of the top-ranked node, indicating that its accessibility to other network nodes and its potential role in modulating signaling are reduced when primary hubs are lost (Figures 3e, 3f, 4d, 4e, and 4f). This observation highlights how perturbations to central nodes can directly affect both receptor-mediated and lipid-mediated pathways, with potential implications for ECS signaling, metabolic regulation, and therapeutic targeting.

Synthetic cannabinoids, including WIN55,212-2 [48], WIN55,212-3 [48], JWH-018 [49], JWH-073 [49], JWH-133 [49], HU-210 [60], HU-211 [60], CP55,940 [50], SR141716A [51], and SR144528 [51], exhibited significant decreases in network metrics following node removal. These effects were most pronounced in Figures 3e and 4e, which depict changes in degree centrality after removing the top one node by betweenness centrality and the top one node by degree centrality, respectively. Several synthetic cannabinoids dropped in degree centrality by nearly two standard deviations, with SR141716A decreasing by over four standard deviations (Figures 3e and 4e). This highlights the critical dependency of these chemicals on central nodes such as CNR1, underscoring how disruption of key receptors can dramatically reduce the functional influence of ligands within the ECS.

Finally, a comparison of eigenvector centrality changes following the removal of the top one versus the top three nodes by degree centrality revealed a dynamic redistribution of influence (Figures 3g, 3h, 4g, and 4h). Removal of a single top node led to substantial decreases in centrality across many nodes, indicating a loss of network cohesion and stability. In contrast, removing the top three nodes produced positive shifts in eigenvector centrality, reflecting a reweighting of influence toward the remaining highly connected and interlinked nodes. This pattern suggests that, in the absence of dominant hubs, the network reorganizes around secondary yet influential nodes, allowing new centers of connectivity to emerge. Such redistribution illustrates the inherent plasticity of the ECS network and its potential for adaptive reconfiguration, maintaining overall structural coherence even when key regulatory nodes are lost.

## Discussion

Our network-based analysis reveals how centrality measures can pinpoint key molecular nodes that facilitate signaling and metabolic integration within the ECS. Across these centrality metrics, distinct patterns emerged. Betweenness and degree centrality highlighted proteins such as CNR1 and TRPV1, reflecting their structural and signaling roles, whereas closeness and eigenvector centrality emphasized chemicals such as CBD, THCV, and CBDV, underscoring their widespread interactions and influence (Figure 2b, Table S12). Together, these findings demonstrate that ECS organization is driven by a set of highly interconnected receptors, enzymes, and ligands that coordinate information flow across receptor-mediated, metabolic, and signaling pathways. The recurring prominence of both canonical receptors (CNR1 and CNR2) and non-canonical components such as GPR55 and TRPV1 underscores the distributed yet integrated nature of ECS signaling. Overall, these centrality-based insights provide a quantitative framework for identifying critical regulatory nodes and potential targets for pharmacological modulation.

Clustering analysis revealed a modular yet interconnected architecture of the ECS network (Figure 2c, Figures S3–S4, Tables S5–S10), in which distinct communities correspond to core biological functions such as receptor-mediated signaling and endocannabinoid metabolism. This modularity reflects the intrinsic organization of the ECS and highlights potential leverage points where pharmacological or genetic perturbations could exert system-wide effects. By linking functional clusters to specific biochemical roles, these findings provide a systems-level framework for investigating how modulation at key nodes might rebalance ECS activity under physiological and pathological conditions.

Our perturbation results also demonstrate that network integrity depends strongly on a subset of highly connected nodes, highlighting the critical influence of nodes in maintaining connectivity and signaling pathways within the ECS. These findings corroborate a previous study [23] that characterized ECS as a scale-free network, in which overall organization and robustness are governed by a relatively small number of highly connected nodes. Perturbation of a significantly important node such as CNR1 reshapes the relative importance of other receptors, enzymes, and ligands, revealing compensatory pathways, as reflected in the increased centrality of CNR2 and GPR55. Concurrently, decrease in network centrality metrics for endocannabinoid-like lipids, such as LEA, illustrate how node loss can reduce accessibility and signaling efficiency, thereby impacting metabolic and receptor-mediated pathways. Understanding the dependencies within these networks offers a framework for predicting how targeting specific receptors, enzymes, or ligands can influence endocannabinoid system signaling, ligand efficacy, or metabolic turnover. This knowledge can inform strategies for drug design, receptor modulation, and combination therapies, which leverage the resilience of the ECS to achieve specific physiological outcomes [23,34,35].

The consistent prominence of GPR55 across multiple network analyses underscores its potential as a key regulatory node within the ECS. In the combined network comprising 80 nodes, GPR55 ranked among the top five nodes for betweenness (5th), closeness (1st), degree (5th), and eigenvector centrality (5th), outperforming other non-canonical GPCRs such as GPR18 and GPR119, which were frequently unranked or placed substantially lower across all measures (Table S13). These trends were even more pronounced in the protein–protein network, which contained 40 nodes, where GPR55 achieved the highest closeness and eigenvector centrality values (both ranked first) as well as high degree centrality (3rd) (Table S13). Its normalized eigenvector centrality score of 0.99 was not only greater than those of GPR18 and GPR119 by 10^15^ and 10^18^ orders of magnitude, respectively, but also the highest observed across all three networks analyzed. High eigenvector centrality values indicate that GPR55 is strongly connected to other highly influential nodes within the ECS, reflecting its central role in maintaining network cohesion and stability. Consistent with this interpretation, perturbation analyses revealed that removal of canonical cannabinoid receptor CNR1 increased GPR55’s relative centrality, suggesting a compensatory or adaptive role in preserving ECS connectivity and signaling integrity. Collectively, these findings position GPR55 as a structurally and functionally significant component of the ECS, supporting growing evidence that it functions as a cannabinoid-responsive receptor involved in modulating synaptic signaling, inflammatory responses, and other physiological processes [40,61,62].

In this study, we constructed a systems-level network of the ECS by integrating experimentally validated interactions between protein and chemical compounds. This network-based framework provides a structural perspective on how key nodes and interactions contribute to ECS signaling (Figures 2a and 2b), how molecular components organize into functional communities (Figure 2c), and how the removal of specific nodes can redirect signaling (Table 1, Figures 3 and 4), reflecting the intrinsic plasticity of the ECS. Although the network captures essential organizational and connectivity patterns, it does not incorporate information on directionality or dynamics. Metrics such as in-degree and out-degree centrality [32], which could reveal the directional flow of signaling, were not included because directional information is not readily available in the public databases used for this study (Figure 1). Similarly, dynamic features such as interaction strength [63–68], temporal responses [69–72], or context-dependent regulation [64,73–76] were not represented, meaning that the current model characterizes structural connectivity rather than functional activity. Despite these limitations, our analysis provides a robust foundation for understanding the ECS organization. Future studies could expand this network to incorporate kinetic parameters, tissue-specific interactions, or experimentally measured signaling dynamics, enabling predictive modeling of how endogenous ligands, phytocannabinoids, and synthetic compounds influence ECS signaling and therapeutic outcomes.

## Conclusion

This study presents a comprehensive systems-level analysis of the ECS by integrating protein–protein and protein–chemical interactions into a combined network. Centrality analyses identified the primary cannabinoid receptors CNR1 and CNR2 as major hubs, reflecting their extensive connectivity with both endogenous and exogenous ligands. In addition to these canonical receptors, other key nodes, including TRPV1 and GPR55, ranked highly across multiple centrality measures, indicating their integrative roles in linking receptor-mediated signaling with metabolic and inflammatory pathways. The prominence of GPR55, in particular, supports emerging evidence [40,61,62] that it functions as a cannabinoid-responsive receptor with broad influence within ECS signaling. Clustering analyses further revealed that key nodes, including the primary cannabinoid receptors CNR1 and CNR2, as well as critical chemical ligands such as the endocannabinoids AEA and 2-AG and various phytocannabinoids, maintain network connectivity and organize into biologically meaningful communities, including receptor-centered and metabolic clusters. Perturbation analysis emphasized the significance of hub nodes in the network. The removal of CNR1 caused substantial disruptions in the network’s organization, reducing the importance of dependent synthetic cannabinoids and shifting centrality to other receptors, such as CNR2 and GPR55, revealing compensatory pathways that support network resilience. These findings demonstrate that the ECS functions as a complex network orchestrating diverse physiological processes, including signaling and metabolic regulation. Characterizing the connectivity of its receptors, enzymes, ligands, and other components provides critical insights for the development of therapeutic strategies and drug design.

## List of abbreviations

ECS: Endocannabinoid system
CNR1: Cannabinoid receptor 1
CNR2: Cannabinoid receptor 2
CB1: Cannabinoid receptor type 1 (alternate name for CNR1)
CB2: Cannabinoid receptor type 2 (alternate name for CNR2)
TRPV1: Transient receptor potential vanilloid 1
GPR55: G-protein coupled receptor 55
GPR18: G-protein coupled receptor 18
GPR119: G-protein coupled receptor 119
GPCR: G-protein coupled receptor
AEA: Anandamide
2-AG: 2-Arachidonoylglycerol
Δ9-THC: Delta-9-tetrahydrocannabinol
THC: Tetrahydrocannabinol
CBD: Cannabidiol
THCV: Tetrahydrocannabivarin
CBDV: Cannabidivarin
OEA: Oleoylethanolamide
LEA: Linoleoyl ethanolamide
FAAH: Fatty acid amide hydrolase
MGLL: Monoacylglycerol lipase
NAPEPLD: N-acyl-phosphatidylethanolamine phospholipase D
DAGLA: Diacylglycerol lipase alpha
DAGLB: Diacylglycerol lipase beta
ABHD6: Abhydrolase domain-containing protein 6
ABHD12: Abhydrolase domain-containing protein 12
FABP7: Fatty acid-binding protein 7
PPARA: Peroxisome proliferator-activated receptor alpha
PTGS2: Prostaglandin-endoperoxide synthase 2 (cyclooxygenase-2)
PLCE1: Phospholipase C epsilon 1
PLCB1: Phospholipase C beta 1
GNAI: Inhibitory Guanine nucleotide-binding protein G alpha subunit
GNAI1: Inhibitory Guanine nucleotide-binding protein G alpha subunit 1
GNAI2: Inhibitory Guanine nucleotide-binding protein G alpha subunit 2
GNAI3: Inhibitory Guanine nucleotide-binding protein G alpha subunit 3
PPI: Protein–protein interaction
PCI: Protein–chemical interaction
SMILES: Simplified Molecular-Input Line-Entry System
IUPAC: International Union of Pure and Applied Chemistry

## Declarations

### Author Contributions

RG, SMD, and AT conceived the project. SMD designed and supervised the project. AS conducted data curation, network construction, computational analyses, and visualizations. AS and SMD analyzed and interpreted the results and wrote and edited the manuscript. AT provided insights on results and contributed to manuscript edits. RG provided insights on the results, contributed manuscript edits, and supported the research through funding.

## Acknowledgements

Authors would like to thank Discovery Partners Institute and the SPARKS internship program for supporting the initiation of this project.

