## Supporting Information for "Integrative Network Analysis Reveals Organizational Principles of the Endocannabinoid System"

**Table S1: Chemical compounds included in the ECS network.** List of chemical nodes (40) with compound names and PubChem CIDs used for integration into the protein–chemical and combined interaction network. Full metadata, including IUPAC names, SMILES strings, and RDKit-calculated descriptors, are provided separately at https://github.com/aanyashridhar/ECS-Network.

| Chemical | PubChem CID |
|---|---|
| AEA | 5281969 |
| 2-AG | 5282280 |
| Noladin | 5311103 |
| Virodhamine | 6443013 |
| NADA | 5311436 |
| AA | 5283678 |
| OEA | 5286420 |
| PEA | 10467 |
| LEA | 5283387 |
| NAGly | 23667530 |
| Δ9-THC | 16078 |
| Δ8-THC | 644019 |
| THCA | 644017 |
| THCV | 644011 |
| CBD | 6440190 |
| CBDA | 644001 |
| CBDV | 92012705 |
| CBG | 5281963 |
| CBGA | 5281962 |
| CBC | 5281965 |
| CBN | 65156 |
| CBL | 5281447 |
| WIN55,212-2 | 5311105 |
| WIN55,212-3 | 5311106 |
| CP55,940 | 443912 |
| HU-210 | 5311352 |
| HU-211 | 146660 |

| | |
|---|---|
| JWH-018 | 588025 |
| JWH-073 | 9850028 |
| JWH-133 | 86287562 |
| AM251 | 6918495 |
| SR141716A | 3003824 |
| SR144528 | 5311135 |
| PGE2 | 5280360 |
| PGD2 | 5280896 |
| PGA2 | 5280895 |
| 12-HETE | 444899 |
| 15-HETE | 5280734 |
| 5-HPETE | 5282820 |
| LTB4 | 5280934 |

**Table S2: Proteins included in the ECS network.** List of protein nodes (40) with protein names and UniProt IDs used for integration into the protein–protein and combined interaction network. Full metadata, including full names, protein types, sub-cellular locations, structural annotations, PDB IDs, amino acid sequences, and evidence codes, are provided separately at https://github.com/aanyashridhar/ECS-Network.

| Protein | UniProt ID |
|---|---|
| CNR1 | P21554 |
| CNR2 | P34972 |
| GPR55 | Q9Y2T6 |
| GPR18 | Q14330 |
| GPR119 | Q8TDV5 |
| TRPV1 | Q8NER1 |
| TRPV2 | Q9Y5S1 |
| TRPV3 | Q8NET8 |
| TRPV4 | Q9HBA0 |
| PPARA | Q07869 |
| PPARD | Q03181 |
| PPARG | P37231 |
| NR1I2 | O75469 |
| NAPEPLD | Q6IQ20 |
| PLCB1 | Q9NQ66 |
| PLCE1 | Q9P212 |
| DAGLA | Q9Y4D2 |
| DAGLB | Q8NCG7 |
| FAAH | O00519 |
| NAAA | Q02083 |
| MGLL | Q99685 |
| ABHD6 | Q9BV23 |
| ABHD12 | Q8N2K0 |
| PTGS2 | P35354 |
| ALOX5 | P09917 |
| ALOX12 | P18054 |
| ALOX15 | P16050 |
| FABP1 | P07148 |

| | |
|---|---|
| FABP3 | P05413 |
| FABP5 | Q01469 |
| FABP7 | O15540 |
| HSPA1A | P0DMV8 |
| SLCO2B1 | O94956 |
| SLC22A5 | O76082 |
| GNAI1 | P63096 |
| GNAI2 | P04899 |
| GNAI3 | P08754 |
| ARRB1 | P49407 |
| GRK2 | P25098 |
| AKT1 | P31749 |

**Table S3: Protein–protein interactions included in the ECS network.** List of protein–protein interactions (92) with interacting protein pairs and confidence scores (thresholded at 0.7) used for integration into the protein–protein and combined interaction network.

| Interacting Protein A | Interacting Protein B | Confidence |
|---|---|---|
| GRK2 | ARRB1 | 0.998 |
| CNR1 | GNAI1 | 0.995 |
| GNAI2 | CNR1 | 0.993 |
| GNAI2 | GNAI3 | 0.992 |
| ARRB1 | AKT1 | 0.99 |
| FABP5 | PPARD | 0.988 |
| GNAI3 | GNAI1 | 0.984 |
| CNR1 | GPR55 | 0.983 |
| CNR1 | CNR2 | 0.981 |
| GNAI2 | GNAI1 | 0.975 |
| FAAH | MGLL | 0.974 |
| PTGS2 | ALOX5 | 0.973 |
| FAAH | CNR1 | 0.972 |
| FABP1 | PPARA | 0.964 |
| ALOX15 | PTGS2 | 0.963 |
| ABHD6 | ABHD12 | 0.949 |
| ALOX12 | PTGS2 | 0.947 |
| MGLL | CNR1 | 0.941 |
| CNR1 | GNAI3 | 0.933 |
| FAAH | NAPEPLD | 0.931 |
| PLCB1 | GNAI3 | 0.929 |
| FABP1 | FABP3 | 0.929 |
| PLCB1 | GNAI1 | 0.928 |
| GNAI2 | PLCB1 | 0.926 |
| GPR119 | GPR55 | 0.922 |
| TRPV1 | GPR55 | 0.922 |
| MGLL | NAPEPLD | 0.92 |
| DAGLA | DAGLB | 0.915 |
| ALOX15 | ALOX5 | 0.914 |
| ALOX12 | ALOX5 | 0.914 |

| | | |
|---|---|---|
| CNR2 | GNAI1 | 0.909 |
| ALOX12 | ALOX15 | 0.906 |
| MGLL | ABHD6 | 0.904 |
| FAAH | DAGLA | 0.898 |
| DAGLA | NAPEPLD | 0.894 |
| FAAH | FABP5 | 0.891 |
| DAGLB | NAPEPLD | 0.89 |
| DAGLA | MGLL | 0.887 |
| PPARD | PPARA | 0.887 |
| FAAH | CNR2 | 0.886 |
| FAAH | ABHD6 | 0.881 |
| GPR119 | GPR18 | 0.868 |
| FAAH | TRPV1 | 0.855 |
| FAAH | NAAA | 0.85 |
| FAAH | GPR55 | 0.848 |
| FAAH | DAGLB | 0.848 |
| MGLL | ABHD12 | 0.839 |
| FAAH | FABP7 | 0.839 |
| FAAH | ABHD12 | 0.831 |
| DAGLA | ABHD6 | 0.831 |
| MGLL | DAGLB | 0.829 |
| PPARG | FABP5 | 0.828 |
| FAAH | FABP3 | 0.826 |
| GPR18 | CNR2 | 0.823 |
| ABHD6 | NAPEPLD | 0.821 |
| DAGLA | CNR1 | 0.817 |
| GPR18 | CNR1 | 0.816 |
| NAAA | NAPEPLD | 0.816 |
| ABHD6 | DAGLB | 0.812 |
| ABHD12 | NAPEPLD | 0.803 |
| NAPEPLD | GPR55 | 0.801 |
| CNR1 | NAPEPLD | 0.8 |
| GPR18 | TRPV1 | 0.796 |
| CNR2 | GPR55 | 0.793 |
| DAGLA | ABHD12 | 0.791 |

| | | |
|---|---|---|
| PPARG | AKT1 | 0.79 |
| CNR1 | TRPV1 | 0.781 |
| MGLL | NAAA | 0.78 |
| GRK2 | AKT1 | 0.778 |
| MGLL | GPR55 | 0.776 |
| PTGS2 | AKT1 | 0.775 |
| TRPV4 | TRPV1 | 0.773 |
| NAPEPLD | TRPV1 | 0.771 |
| GPR18 | GPR55 | 0.767 |
| CNR1 | ARRB1 | 0.757 |
| DAGLB | ABHD12 | 0.746 |
| PPARA | GPR55 | 0.746 |
| ARRB1 | GPR55 | 0.744 |
| FAAH | GPR18 | 0.736 |
| PPARG | PTGS2 | 0.735 |
| PTGS2 | PPARA | 0.729 |
| MGLL | TRPV1 | 0.728 |
| PPARA | TRPV1 | 0.718 |
| ABHD6 | GPR55 | 0.717 |
| PPARG | PPARD | 0.715 |
| FABP5 | PPARA | 0.708 |
| GPR119 | PPARA | 0.706 |
| DAGLB | CNR1 | 0.705 |
| PLCB1 | PLCE1 | 0.705 |
| NAAA | PPARA | 0.702 |
| PPARA | AKT1 | 0.7 |
| DAGLA | GPR55 | 0.7 |

**Table S4: Protein–chemical interactions included in the ECS network.** List of protein–chemical interactions (126) with interacting chemicals and interacting proteins used for integration into the protein–chemical and combined interaction network.

| Chemical | Protein |
|---|---|
| AEA | CNR1 |
| AEA | CNR2 |
| AEA | FAAH |
| AEA | MGLL |
| AEA | TRPV1 |
| 2-AG | MGLL |
| 2-AG | CNR2 |
| 2-AG | CNR1 |
| 2-AG | DAGLB |
| 2-AG | DAGLA |
| 2-AG | GPR119 |
| 2-AG | FAAH |
| NADA | TRPV1 |
| NADA | FAAH |
| AA | PTGS2 |
| AA | ALOX5 |
| OEA | GPR119 |
| OEA | NAAA |
| OEA | CNR2 |
| OEA | PPARA |
| OEA | TRPV1 |
| OEA | PPARD |
| OEA | CNR1 |
| PEA | NAAA |
| LEA | TRPV1 |
| LEA | CNR1 |
| NAGly | TRPV1 |
| NAGly | FAAH |
| Δ9-THC | GPR18 |
| Δ9-THC | PPARD |

| | |
|---|---|
| Δ8-THC | GPR18 |
| Δ8-THC | PPARD |
| THCA | ALOX15 |
| THCV | CNR2 |
| THCV | TRPV1 |
| THCV | TRPV2 |
| THCV | TRPV4 |
| THCV | CNR1 |
| THCV | NAAA |
| THCV | GPR55 |
| THCV | TRPV3 |
| THCV | DAGLA |
| CBD | PPARG |
| CBD | NR1I2 |
| CBD | CNR1 |
| CBD | TRPV1 |
| CBD | PTGS2 |
| CBD | CNR2 |
| CBD | DAGLA |
| CBD | NAAA |
| CBD | GPR55 |
| CBDA | ALOX15 |
| CBDV | GPR55 |
| CBDV | CNR2 |
| CBDV | TRPV2 |
| CBDV | DAGLA |
| CBDV | TRPV1 |
| CBDV | TRPV4 |
| CBDV | NAAA |
| CBDV | TRPV3 |
| CBDV | CNR1 |
| CBG | PPARG |
| CBG | TRPV2 |
| CBG | TRPV1 |
| CBG | CNR2 |

| | |
|---|---|
| CBG | CNR1 |
| CBG | DAGLA |
| CBG | TRPV3 |
| CBG | NAAA |
| CBG | TRPV4 |
| CBGA | DAGLA |
| CBGA | NAAA |
| CBGA | CNR1 |
| CBGA | ALOX15 |
| CBC | TRPV3 |
| CBC | TRPV4 |
| CBC | CNR2 |
| CBC | DAGLA |
| CBC | NAAA |
| CBC | TRPV2 |
| CBC | PPARG |
| CBC | TRPV1 |
| CBC | CNR1 |
| CBN | CNR2 |
| CBN | CNR1 |
| CBN | TRPV1 |
| CBN | TRPV2 |
| CBN | NAAA |
| CBN | TRPV4 |
| CBN | DAGLA |
| CBN | PPARG |
| CBN | TRPV3 |
| WIN55,212-2 | CNR1 |
| WIN55,212-2 | CNR2 |
| WIN55,212-3 | CNR1 |
| WIN55,212-3 | CNR2 |
| CP55,940 | CNR1 |
| CP55,940 | CNR2 |
| HU-210 | CNR2 |
| HU-210 | CNR1 |

| | |
|---|---|
| HU-211 | CNR1 |
| HU-211 | CNR2 |
| JWH-018 | CNR2 |
| JWH-018 | CNR1 |
| JWH-073 | CNR2 |
| JWH-133 | CNR2 |
| JWH-133 | CNR1 |
| JWH-133 | TRPV1 |
| AM251 | CNR1 |
| AM251 | TRPV1 |
| AM251 | GPR55 |
| AM251 | CNR2 |
| SR141716A | CNR1 |
| SR144528 | CNR2 |
| SR144528 | CNR1 |
| PGE2 | PTGS2 |
| PGD2 | PPARG |
| PGD2 | PPARA |
| PGA2 | PTGS2 |
| 12-HETE | ALOX5 |
| 12-HETE | ALOX15 |
| 12-HETE | ALOX12 |
| 12-HETE | PTGS2 |
| 12-HETE | PPARA |
| 15-HETE | ALOX5 |
| LTB4 | ALOX5 |

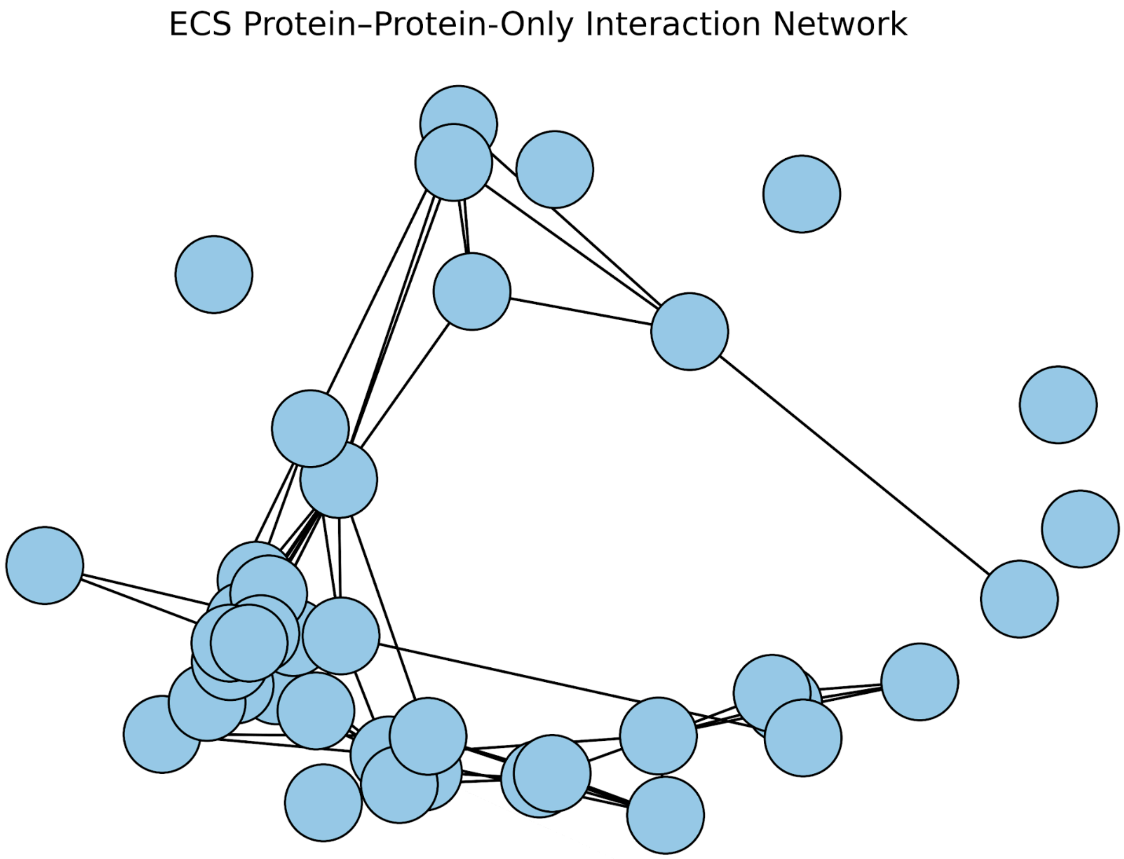


**Figure S1: Protein–protein interaction network.** Visualization of the protein–protein only network showing proteins connected through documented interactions. The network has 40 nodes and 92 edges.

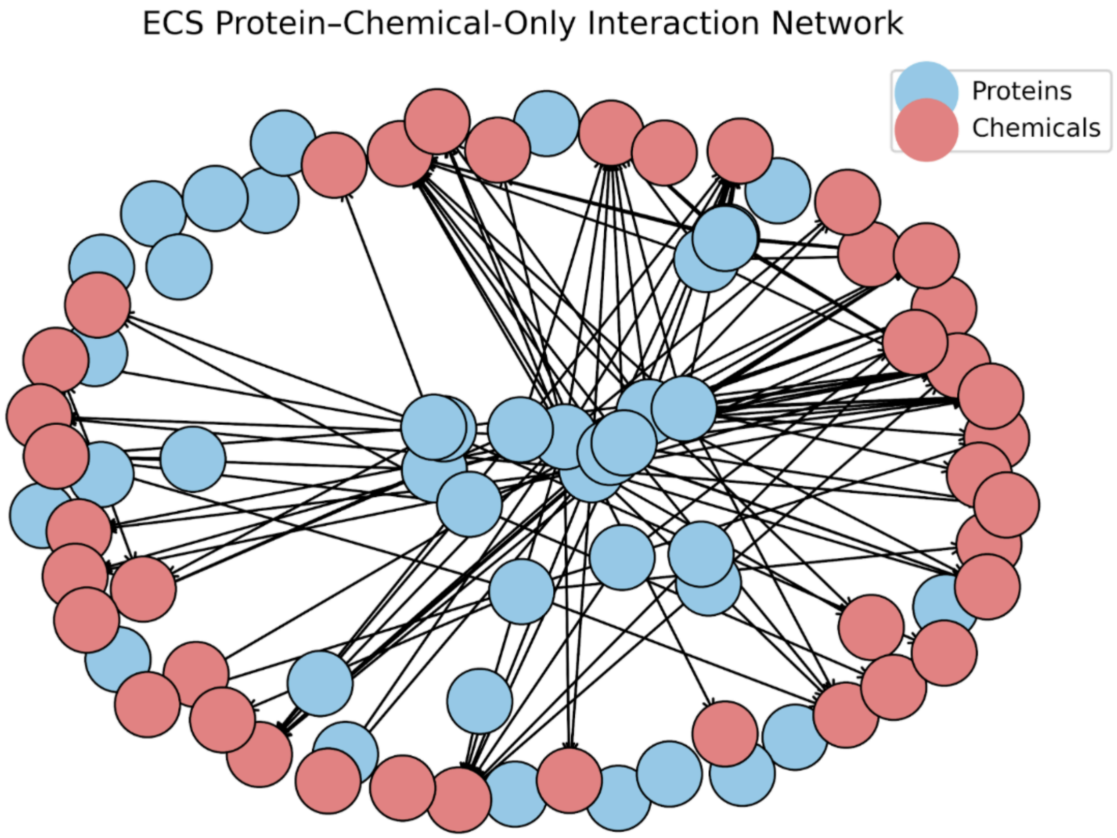


**Figure S2: Protein–chemical interaction network.** Visualization of the protein–chemical only network (with protein–protein interactions filtered out) showing proteins (blue) and chemicals (red) connected through documented interactions. The network has 40 nodes and 126 edges.

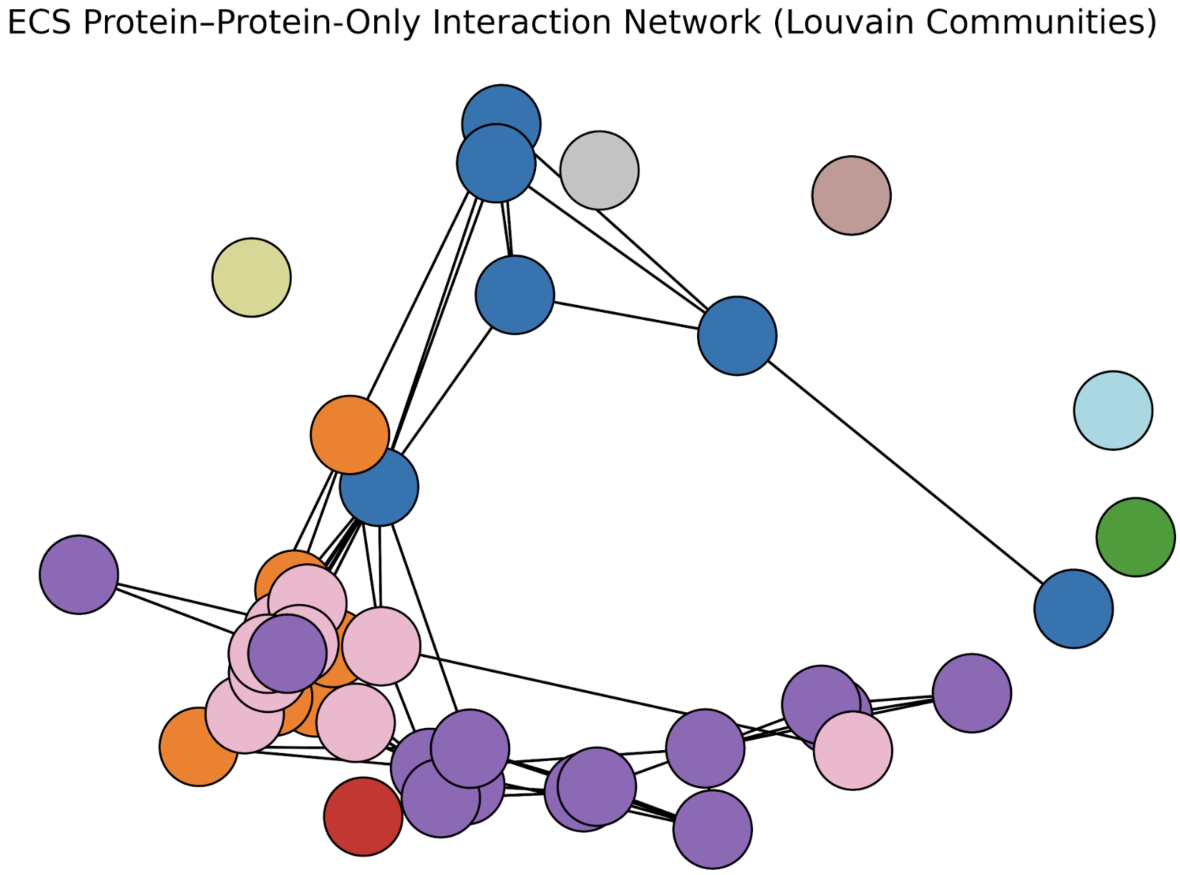


**Figure S3: Protein–protein interaction network Louvain communities from clustering analysis.** Louvain community detection applied to the protein–protein only network, partitioning the nodes into modular clusters (denoted by distinct colors) based on interaction density, which highlights potential functional or mechanistic groupings within ECS proteins.

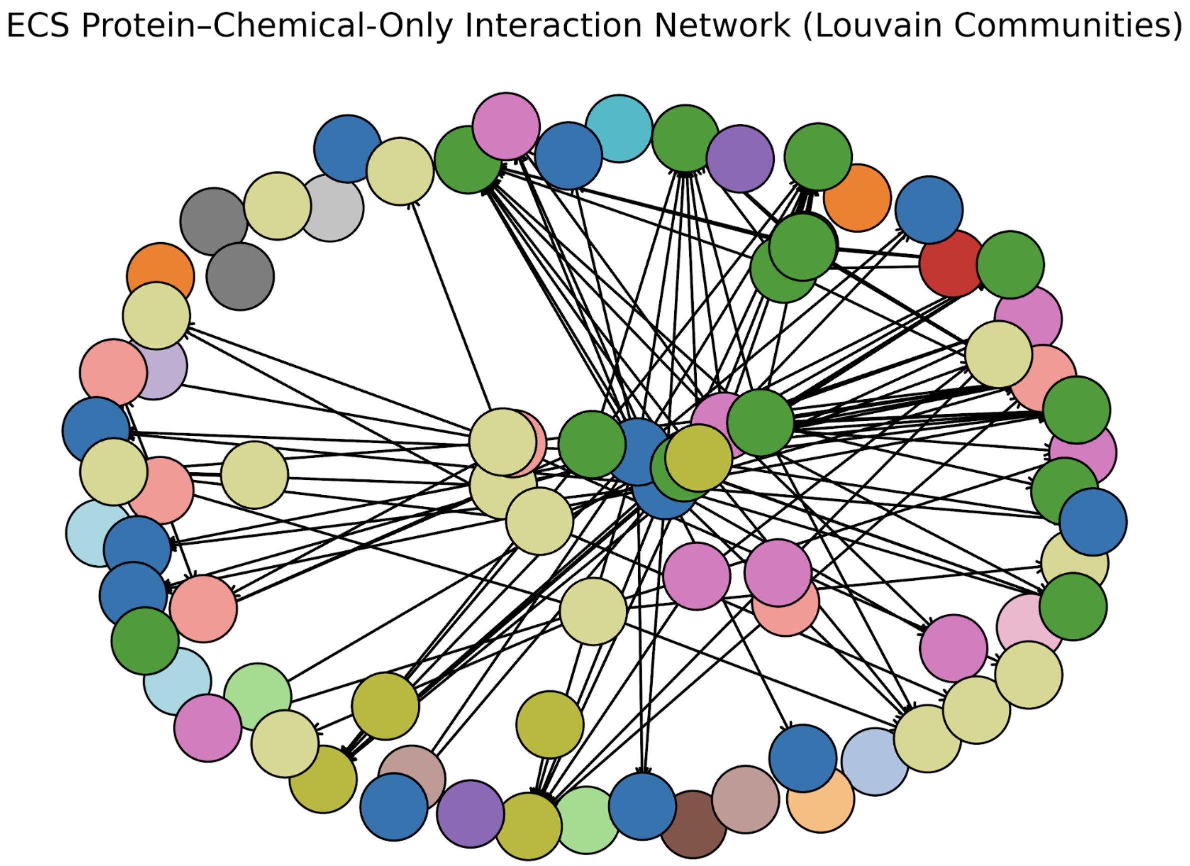


**Figure S4: Protein–chemical interaction network Louvain communities from clustering analysis.** Louvain community detection applied to the protein–chemical only network (with protein–protein interactions filtered out), partitioning the nodes into modular clusters (denoted by distinct colors) based on interaction density, which highlights potential functional or mechanistic groupings within ECS proteins and chemicals.

**Table S5: Louvain community assignments for the combined interaction network.** List of proteins and chemicals and their assigned Louvain communities.

| Node | Louvain Community |
|---|---|
| CNR1 | 0 |
| CNR2 | 0 |
| PLCE1 | 0 |
| PLCB1 | 0 |
| AEA | 0 |
| GNAI3 | 0 |
| GNAI1 | 0 |
| GNAI2 | 0 |
| WIN55,212-3 | 0 |
| WIN55,212-2 | 0 |
| JWH-133 | 0 |
| JWH-073 | 0 |
| JWH-018 | 0 |
| HU-211 | 0 |
| HU-210 | 0 |
| CP55,940 | 0 |
| SR141716A | 0 |
| AM251 | 0 |
| SR144528 | 0 |
| SLCO2B1 | 1 |
| FAAH | 2 |
| NAPEPLD | 2 |
| DAGLB | 2 |
| ABHD12 | 2 |
| 2-AG | 2 |
| FABP7 | 2 |
| ABHD6 | 2 |
| MGLL | 2 |
| GPR55 | 2 |
| DAGLA | 2 |
| SLC22A5 | 3 |

| | |
|---|---|
| THCV | 4 |
| CBC | 4 |
| CBN | 4 |
| CBDV | 4 |
| PEA | 4 |
| CBG | 4 |
| TRPV2 | 4 |
| TRPV3 | 4 |
| NADA | 4 |
| TRPV4 | 4 |
| NAAA | 4 |
| LEA | 4 |
| NAGly | 4 |
| TRPV1 | 4 |
| GRK2 | 5 |
| Δ8-THC | 5 |
| ARRB1 | 5 |
| AKT1 | 5 |
| OEA | 5 |
| GPR119 | 5 |
| GPR18 | 5 |
| PPARA | 5 |
| FABP3 | 5 |
| PGD2 | 5 |
| Δ9-THC | 5 |
| PPARG | 5 |
| PPARD | 5 |
| FABP1 | 5 |
| FABP5 | 5 |
| Noladin | 6 |
| CBD | 7 |
| NR1I2 | 7 |
| PTGS2 | 8 |
| ALOX15 | 8 |
| ALOX12 | 8 |

| | |
|---|---|
| PGE2 | 8 |
| THCA | 8 |
| AA | 8 |
| CBDA | 8 |
| ALOX5 | 8 |
| CBGA | 8 |
| 12-HETE | 8 |
| 15-HETE | 8 |
| LTB4 | 8 |
| PGA2 | 8 |
| Virodhamine | 9 |
| CBL | 10 |
| 5-HPETE | 11 |
| HSPA1A | 12 |

**Table S6: Louvain community summary metrics for the combined interaction network.** Report of overall network modularity, the total number of detected communities, and community size statistics.

| Metric | Value |
|---|---|
| Overall Modularity | 0.4277 |
| Number of Communities | 13 |
| Largest Community Size | 19 |
| Smallest Community Size | 1 |

**Table S7: Louvain community assignments for the protein–protein interaction network.** List of proteins and their assigned Louvain communities.

| Protein | Louvain Community |
|---|---|
| CNR1 | 0 |
| PLCB1 | 0 |
| PLCE1 | 0 |
| GNAI2 | 0 |
| GNAI1 | 0 |
| GNAI3 | 0 |
| GPR119 | 1 |
| CNR2 | 1 |
| TRPV4 | 1 |
| TRPV1 | 1 |
| GPR55 | 1 |
| GPR18 | 1 |
| TRPV2 | 2 |
| TRPV3 | 3 |
| PPARG | 4 |
| PPARA | 4 |
| FABP1 | 4 |
| ALOX5 | 4 |
| FABP5 | 4 |
| FABP3 | 4 |
| ALOX12 | 4 |
| PPARD | 4 |
| ALOX15 | 4 |
| PTGS2 | 4 |
| ARRB1 | 4 |
| GRK2 | 4 |
| AKT1 | 4 |
| NR1I2 | 5 |
| ABHD12 | 6 |
| ABHD6 | 6 |
| NAPEPLD | 6 |

| | |
|---|---|
| DAGLA | 6 |
| NAAA | 6 |
| FABP7 | 6 |
| FAAH | 6 |
| DAGLB | 6 |
| MGLL | 6 |
| SLC22A5 | 7 |
| SLCO2B1 | 8 |
| HSPA1A | 9 |

**Table S8: Louvain community summary metrics for the protein–protein interaction network.** Report of overall network modularity, the total number of detected communities, and community size statistics.

| Metric | Value |
|---|---|
| Overall Modularity | 0.4207 |
| Number of Communities | 10 |
| Largest Community Size | 13 |
| Smallest Community Size | 1 |

**Table S9: Louvain community assignments for the protein–chemical interaction network.** List of proteins and chemicals and their assigned Louvain communities.

| Node | Louvain Community |
|---|---|
| CNR1 | 0 |
| CNR2 | 0 |
| WIN55,212-3 | 0 |
| WIN55,212-2 | 0 |
| JWH-018 | 0 |
| HU-211 | 0 |
| HU-210 | 0 |
| CP55,940 | 0 |
| SR141716A | 0 |
| SR144528 | 0 |
| JWH-073 | 0 |
| GNAI1 | 1 |
| GNAI2 | 2 |
| GNAI3 | 3 |
| ARRB1 | 4 |
| GRK2 | 5 |
| CBG | 6 |
| CBN | 6 |
| CBC | 6 |
| THCV | 6 |
| CBDV | 6 |
| TRPV4 | 6 |
| PPARG | 6 |
| PGD2 | 6 |
| AM251 | 6 |
| PEA | 6 |
| GPR55 | 6 |
| TRPV3 | 6 |
| TRPV2 | 6 |
| NAAA | 6 |
| AKT1 | 7 |

| | |
|---|---|
| Noladin | 8 |
| Virodhamine | 9 |
| PPARD | 10 |
| GPR18 | 10 |
| GPR119 | 10 |
| Δ8-THC | 10 |
| OEA | 10 |
| Δ9-THC | 10 |
| CBL | 11 |
| 5-HPETE | 12 |
| NAPEPLD | 13 |
| PLCB1 | 14 |
| PLCE1 | 15 |
| FABP7 | 16 |
| NADA | 17 |
| AEA | 17 |
| TRPV1 | 17 |
| FAAH | 17 |
| MGLL | 17 |
| NAGly | 17 |
| LEA | 17 |
| JWH-133 | 17 |
| HSPA1A | 18 |
| SLCO2B1 | 19 |
| ABHD6 | 20 |
| ABHD12 | 21 |
| DAGLB | 22 |
| 2-AG | 22 |
| DAGLA | 22 |
| NR1I2 | 22 |
| CBD | 22 |
| ALOX12 | 23 |
| ALOX15 | 23 |
| PPARA | 23 |
| PTGS2 | 23 |

| | |
|---|---|
| ALOX5 | 23 |
| PGE2 | 23 |
| AA | 23 |
| CBGA | 23 |
| THCA | 23 |
| CBDA | 23 |
| PGA2 | 23 |
| 12-HETE | 23 |
| 15-HETE | 23 |
| LTB4 | 23 |
| SLC22A5 | 24 |
| FABP1 | 25 |
| FABP3 | 26 |
| FABP5 | 27 |

**Table S10: Louvain community summary metrics for the protein–chemical interaction network.** Report of overall network modularity, the total number of detected communities, and community size statistics.

| Metric | Value |
|---|---|
| Overall Modularity | 0.3969 |
| Number of Communities | 28 |
| Largest Community Size | 14 |
| Smallest Community Size | 1 |

**Table S11: Protein–protein interaction network centrality metrics.** Top five nodes ranked by centrality measures in the protein–protein only network. Full rankings and centrality metric calculations can be found at https://github.com/aanyashridhar/ECS-Network.

| | Betweenness | Closeness | Degree | Eigenvector |
|---|---|---|---|---|
| 1 | CNR1 | GPR55 | FAAH | GPR55 |
| 2 | PPARA | TRPV1 | CNR1 | TRPV1 |
| 3 | PTGS2 | AKT1 | GPR55 | GNAI1 |
| 4 | ARRB1 | NAPEPLD | NAPEPLD | AKT1 |
| 5 | GPR18 | PPARA | MGLL | NAPEPLD |

**Table S12: Protein–chemical interaction network centrality metrics.** Top five nodes ranked by centrality measures in the protein–chemical only network. Betweenness centrality is excluded in the protein–chemical network as this centrality relies on shortest path calculations between nodes, which are not well-defined in networks where chemicals only connect to proteins. Full rankings and centrality metric calculations can be found at https://github.com/aanyashridhar/ECS-Network.

| | Closeness | Degree | Eigenvector |
|---|---|---|---|
| 1 | CBN | CNR1 | CBN |
| 2 | CBC | CNR2 | CBC |
| 3 | CBD | TRPV1 | CBD |
| 4 | THCV | CBDV | THCV |
| 5 | CBG | CBN | CBG |

**Table S13: Comparative centrality rankings of G-protein coupled proteins, GPR55, GPR18, and GPR119.** Rankings of three non-canonical cannabinoid receptors across the combined ECS network and the protein–protein only network showing GPR55 outperforming GPR18 and GPR119 in nearly every metric, with the exception of betweenness centrality in the protein–protein only network. Unranked indicates a centrality value of zero, meaning the node had no measurable contribution for that metric. Full rankings and centrality metric calculations can be found at https://github.com/aanyashridhar/ECS-Network.

| Network | Centrality | **GPR55** | **GPR18** | **GPR119** |
|---|---|---|---|---|
| Combined | Betweenness | 5 | 8 | Unranked |
| Combined | Closeness | 1 | 51 | Unranked |
| Combined | Degree | 5 | 19 | 29 |
| Combined | Eigenvector | 5 | 56 | 63 |
| Protein–Protein | Betweenness | Unranked | 5 | Unranked |
| Protein–Protein | Closeness | 1 | 18 | Unranked |
| Protein–Protein | Degree | 3 | 11 | 24 |
| Protein–Protein | Eigenvector | 1 | 20 | 27 |
